# The phylogenetic position of the extinct Hawaiian honeyeaters: Overcoming the limitations of antique DNA

**DOI:** 10.1101/2025.06.02.657453

**Authors:** Min Zhao, Rebecca T. Kimball, Edward L. Braun

**Affiliations:** Department of Biology, University of Florida, Gainesville, FL 32611, US; Department of Biological Sciences, Virginia Tech, Blacksburg, VA 24061, US

## Abstract

Mohoidae, the Hawaiian honeyeaters, are a group of extinct birds that once inhabited the Hawaiian Islands. Their genetic data are only available from historical museum specimens, so it is based on highly fragmented and damaged DNA. Poor sequence recovery in these “antique” samples can bias phylogenetic inference which often results in exceptionally long terminal branches in the phylogeny. These long branches can distort estimates of phylogeny, an artifact known as long branch attraction. However, the best way to analyze this data is unclear because these long branches are typically artifactual, reflecting DNA quality. We investigated the position of Mohoidae within the superfamily Bombycilloidea using published alignments of ultraconserved elements (UCEs) and re-assembled UCEs from raw sequencing reads. The uncertainty involves two monotypic families (Hypocoliidae and Hylocitreidae). Re-aligning UCEs using a smaller taxon set closely related to Mohoidae, along with edge trimming and re-assembly, eliminated the long branches of Mohoidae and Hypocoliidae (also represented by an antique sample). The sister relationship of these two families in previous studies appears to be artifactual; our new analyses yielded congruent topologies that placed Mohoidae as sister to a clade comprising Hypocoliidae and Hylocitreidae. We also found that different assemblers (SPAdes vs Trinity) yielded sequences with conflicts towards the ends of the UCEs, especially for the antique samples, although this did not alter the topology. Finally, combining the UCEs with legacy data allowed us to produce a strongly supported complete phylogeny of Bombycilloidea the included all extant species as well as the extinct Mohoidae.

## Introduction

Understanding the evolutionary history of extinct species is critical for reconstructing a complete and reliable Tree of Life. It provides a more accurate picture of how different lineages are related and how biodiversity has changed over time. However, while DNA can survive hundreds or thousands of years, ancient DNA samples typically suffer from degradation, such as reduced fragment size, depurination and cytosine deamination (Briggs et al., 2007; Dabney et al., 2013; Pääbo et al., 2004). DNA extracted from more recent museum specimens and other types of natural history collections (“antique DNA”) is typically of higher quality than ancient DNA extracted from fossils (Wandeler et al., 2007), although it is often associated with reduced fragment sizes and contains varying levels of hydrolytic and oxidative damage depending on the preservation methods, storage conditions, and age of the specimen (Wandeler et al., 2003; Zimmermann et al., 2008).

Poor sequence recovery can bias phylogenetic relationships. When sequences are missing or contain errors, they can result in erroneous alignments and incorrect estimates of genetic divergences among taxa, leading to distorted topology and biased branch length estimates (e.g., Hosner et al., 2016; Philippe et al., 2011; Roure et al., 2013). In some cases, taxa with poor sequence recovery appear to have exceptionally long terminal branches (e.g., Bernstein and Ruane, 2022; Braun et al., 2024; Derkarabetian et al., 2019; Kimball et al., 2021; McCullough et al., 2024; Nash et al., 2024; Zhao et al., 2025a). While long branches can result from accelerated sequence evolution (e.g., Braun et al., 2019; Cai et al., 2019; James et al., 2013), those associated with poor data recovery are likely to be methodological artifacts rather than true evolutionary signals (Braun et al., 2024; Kimball et al., 2021; McCullough et al., 2024; Zhao et al., 2025a). The incorrect grouping of two or more long branches is an artifact called long branch attraction (reviewed by Bergsten, 2005) and a major goal of efforts to improve models of sequence evolution has been the reduction of artifacts like long branch attraction (e.g., Kapli et al., 2020; Philippe et al., 2011; Smith et al., 2015). However, the patterns in the data that lead to long branches in analyses including antique samples (where sequencing errors and missing data contribute to terminal branch length) are likely to differ from the site patterns generated by accelerated evolution. Taxa with sequencing errors and extensive missing data may have an even larger impact on multispecies coalescent methods based on combining estimated gene trees because the problematic sequences are non-randomly distributed (Xi et al., 2016). Overall, artifactual long branches might lead to the incorrect placement of taxa that cannot be addressed using standard phylogenetic methods; unfortunately, this possibility has not been explored extensively (but see Xia et al., 2003).

If sequencing errors can be a source of long branches, the choice of bioinformatic tools used to process raw sequence data might have a large impact on downstream alignment and analyses. These choices include the sequence assembler (e.g., Islam et al., 2021; Pightling et al., 2015; Ribeiro et al., 2021), alignment programs (e.g., Pais et al., 2014; Portik and Wiens, 2021; Thompson et al., 2011), and the choice of reference genome if one is used (e.g., Rick et al., 2024; Valiente-Mullor et al., 2021). Assemblers are less likely to be biased when they are used with good quality sequence data with high coverage (Pightling et al., 2014). But the performance of specific assemblers is likely to become more important when the sequencing depth is more variable, which is expected to be the case for sequence capture data (Faircloth et al., 2012), transcriptome sequencing (Hölzer and Marz, 2019), and single-cell sequencing (Daley and Smith, 2014). Choices among bioinformatic pipelines may become even more important when antique samples are analyzed because factors such as DNA fragmentation and damage are likely to further exacerbate these problems. The influence of sequence assembler choice on phylogenomic studies has received limited attention, since many phylogenomic studies focus on using one assembler for all data.

Multiple initiatives across the globe have begun to generate genomic data for many parts of the Tree of Life, such as the Vertebrate Genomes Project (VGP), the Bird 10,000 Genomes (B10K) Project, the OpenWings Project, the 5,000 Insect Genomes Initiative (i5K) and the Ten Thousand Plant Genome Project (10KP). Efforts to build a complete Tree of Life will no doubt use data generated by these initiatives. While it might be viewed as a best practice to begin with raw sequence and perform all analyses including sequence assembly using a single pipeline, it is more typical to use existing assemblies. For well-studied groups like birds there are many ongoing efforts to generate sequence data using multiple methods, such as whole genome sequencing (WGS) (Jarvis et al., 2014; Lamichhaney et al., 2015; Sackton et al., 2019; Stiller et al., 2024), sequence capture (Harvey et al., 2020; McCormack et al., 2013; Moyle et al., 2016; Smith et al., 2023), and transcriptome sequencing (Kuhl et al. 2021). Sequence capture is likely to be especially important for groups that can only be obtained as museum specimens (McCormack et al., 2016; McCullough et al., 2019; Zhao et al., 2025a). In these cases, it is not always clear how integrating these types of heterogeneous sequence data might affect estimates of phylogeny.

The extinct Hawaiian honeyeaters (Mohoidae) were a group of birds that specialized on nectar feeding (Billerman, 2020). Mohoidae includes at least five species from two genera, *Moho* and *Chaetoptila*, that went extinct in the past two centuries (Lovette, 2008); in addition to the historical records of Mohoidae there is fossil evidence for one or two additional *Chaetoptila* species (James and Olson, 1991). While it is clear that Mohoidae fits within the passerines or perching birds (Passeriformes, the largest avian order), its placement has been variable. Mohoidae were originally treated as Australasian honeyeaters (Meliphagidae) due to morphological similarities that reflect their adaptation for nectar-feeding, as well as plumage and biogeographical distribution pattern (Lovette, 2008). However, molecular data (Fleischer et al., 2008) revealed that the Hawaiian honeyeaters were most closely related with three small families: Bombycillidae (waxwings), Ptiliogonatidae (silky flycatchers), and monotypic Dulidae (Palmchat). Other studies (Alström et al., 2014; Braun et al., 2024; Burleigh et al., 2015; Oliveros et al., 2019; Spellman et al., 2008; Zhao et al., 2025b) subsequently found evidence for a clade, which Alström et al. (2014) named Bombycilloidea, comprising Mohoidae, Bombycillidae, Ptiliogonatidae, and Dulidae, as well as two other monotypic families, Hylocitreidae (Hylocitrea) and Hypocoliidae (Gray Hypocolius).

Two previous phylogenomic studies, Oliveros et al., (2019) and Braun et al., (2024), sampled representatives of all six families in Bombycilloidea using thousands of ultraconserved elements (UCEs). Both were large-scale studies, with the former study focused across all passerines, whereas the latter included representatives for almost all avian families. The Mohoidae and Hypocoliidae sequences in these two studies were generated from toe pads cut of antique study skins collected in 1901 and 1937, respectively; sequences for the other four families were generated from frozen tissues. Oliveros et al. (2019) used one assembler (SPAdes) for toe pads and a different assembler (Trinity) for tissues; Braun et al. (2024) used the Oliveros et al. (2019) assemblies for these taxa, though these data were re-aligned. In Braun et al. (2024), both *Moho* and *Hypocolius* were represented by exceptionally long terminal branches and connected by a relatively short internal branch (see also Fig. 1), raising the concern that the sister relationship could reflect long branch attraction. To address this concern, we asked several questions. First, we asked how choice of assemblers might affect phylogenetic inference when incorporating sequences from antique samples. Second, we asked whether adding taxa might aid in breaking up the long branches. Third, we asked whether re-aligning loci using a smaller, more focused taxon set might resolve phylogenetic noise introduced by missing data and potential alignment errors. Lastly, we provided a comprehensive, well-resolved phylogeny for the Bombycilloidea superfamily by combining UCEs with legacy loci (mitochondrial and nuclear loci) that have been commonly used in many previous phylogenetic studies.

**Fig. 1.**
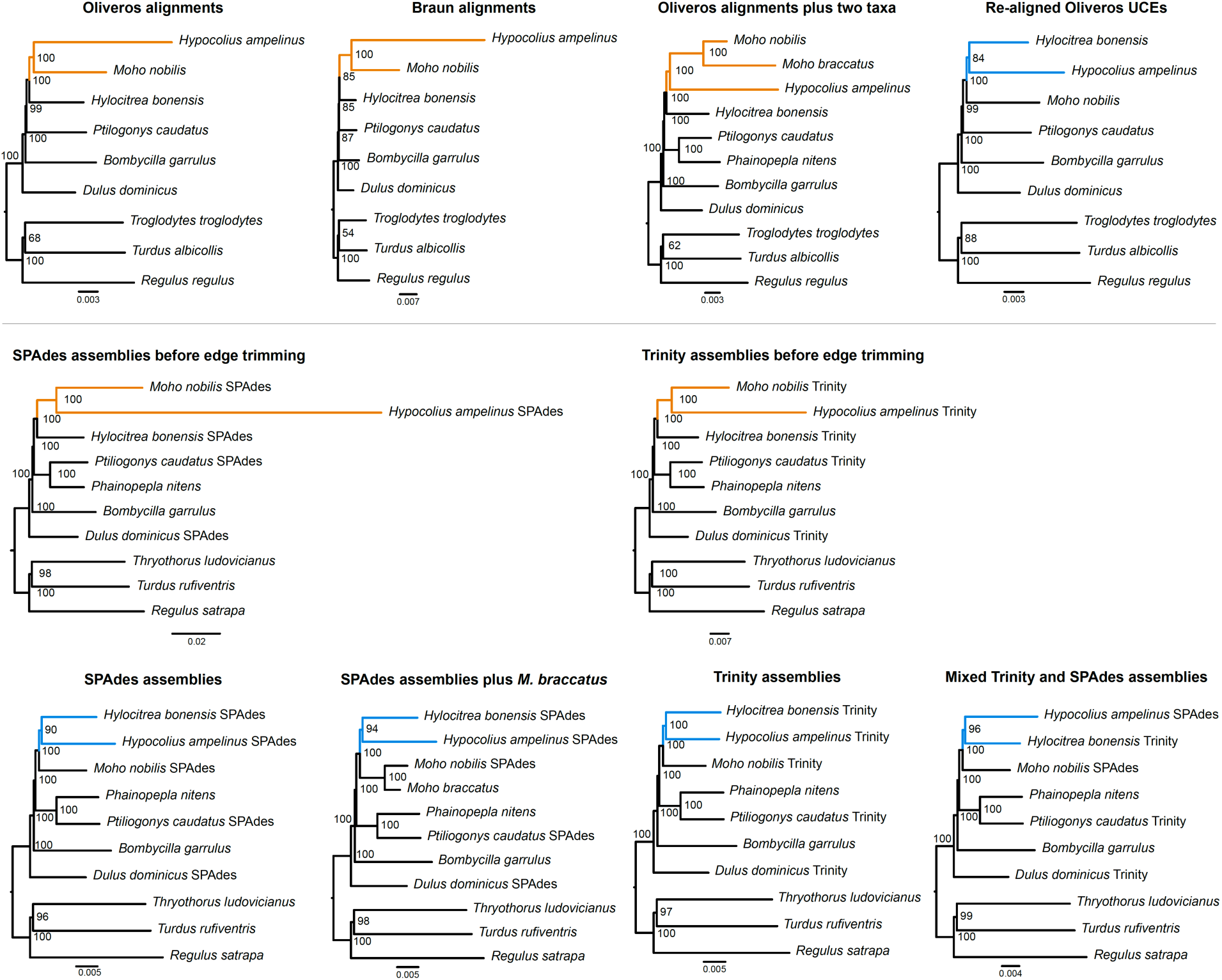
Maximum likelihood trees estimated based on 10 UCE datasets (re-assembled datasets below black line). Except for slight differences in taxon sampling, their topologies only differ in the arrangements among *Moho*, *Hypocolius* and *Hylocitrea* (marked in orange and blue: orange – *Hypocolius* is sister to *Moho*; blue – *Hypocolius* is sister to *Hylocitrea*). Values at nodes are ultrafast bootstrap support values based on 1000 replicates.

## Methods

### Estimating trees using published alignments

We downloaded the complete UCE alignments directly from Oliveros et al. (2019) and Braun et al. (2024) and extracted sequences only for the six Bombycilloidea species and three outgroup species (Table 1) from the supermatrices; referred as Oliveros alignment and Braun alignment respectively. Oliveros et al. (2019) used two different assemblers: Trinity (Hölzer and Marz, 2019) was used for fresh tissue, while SPAdes (Prjibelski et al., 2020) was used for antique (toe pad) samples. Braun et al. (2024) used the Oliveros et al. (2019) UCE assemblies, but then added other species (distantly related taxa that are not part of this study) and re-aligned the data. In addition, internal trimming was used on the Braun alignment, but not on the Oliveros alignment. Both studies used MAFFT (Katoh and Standley, 2013) as the sequence aligner. Three outgroup species were chosen to represent Muscicapoidea, Regulidae and Certhioidea. Though the exact relationships among these outgroups and Bombycilloidea have not been fully resolved (cf. Braun et al., 2024; Oliveros et al., 2019; Stiller et al., 2024; Zhao et al., 2025b), it is clear that Bombycilloidea is most closely related to these three clades. Therefore, we included a total of nine species in the subset alignments. Since these alignments came from datasets with many more taxa, we then used a custom perl script to remove any all-gap (“N”, “?” and “-”) columns in these alignments (this step may not be necessary as most tree inference programs such as IQ-TREE would ignore those columns). For each dataset (Table 2; data sets 1-2), we used IQ-TREE2 v.2.2.2 (Minh et al., 2020) to run a concatenated analysis using the implemented ModelFinder (Kalyaanamoorthy et al., 2017) to select the best-fit partitioning models (-m TESTMERGE) while allowing each partition to have its own rate (-p) and reducing the computational burden with a relaxed hierarchical clustering algorithm (--rcluster 10). Branch support was assessed using 1000 ultrafast bootstrap replicates (-B 1000) (Hoang et al., 2018).

**Table 1.**
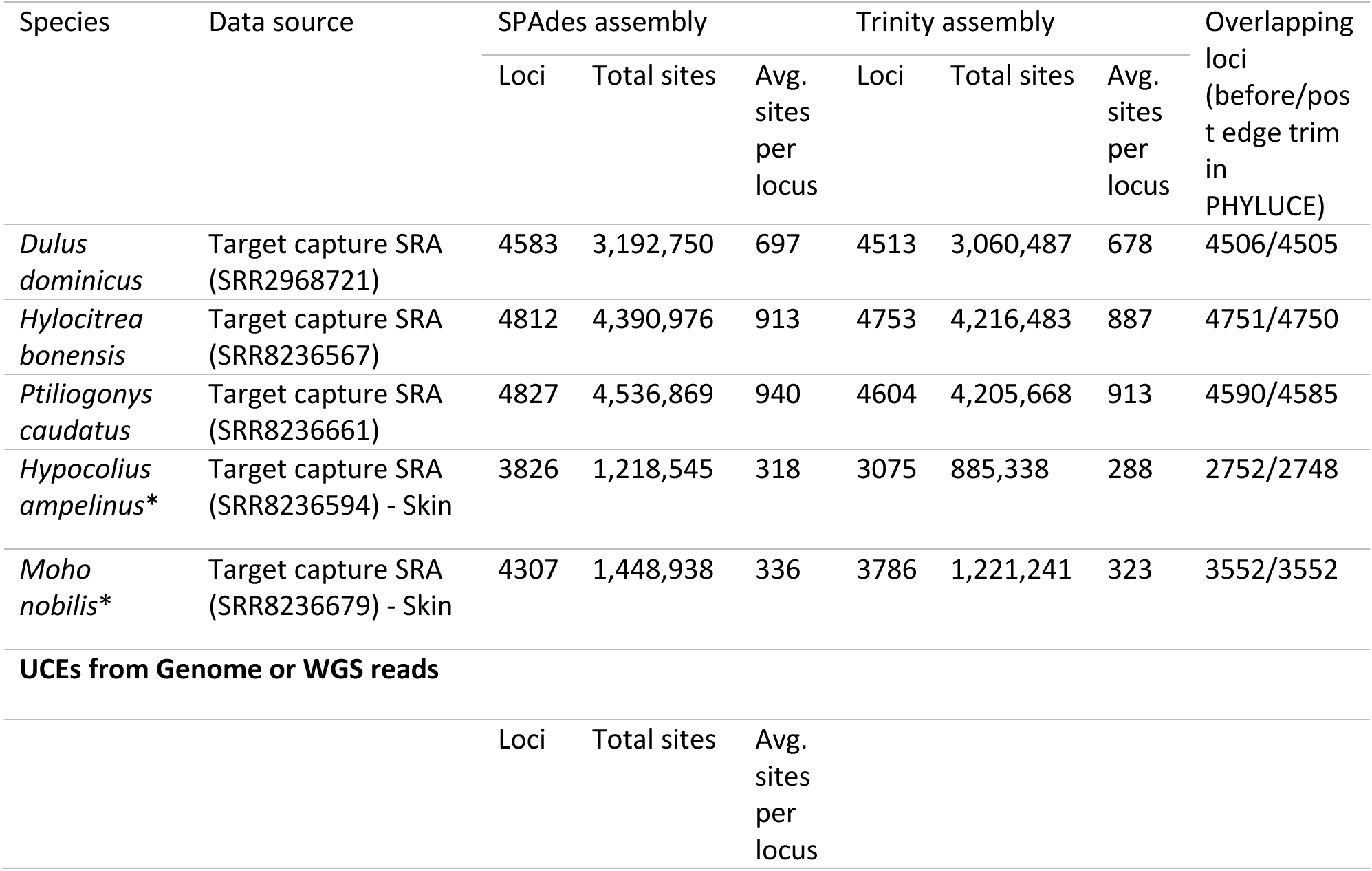
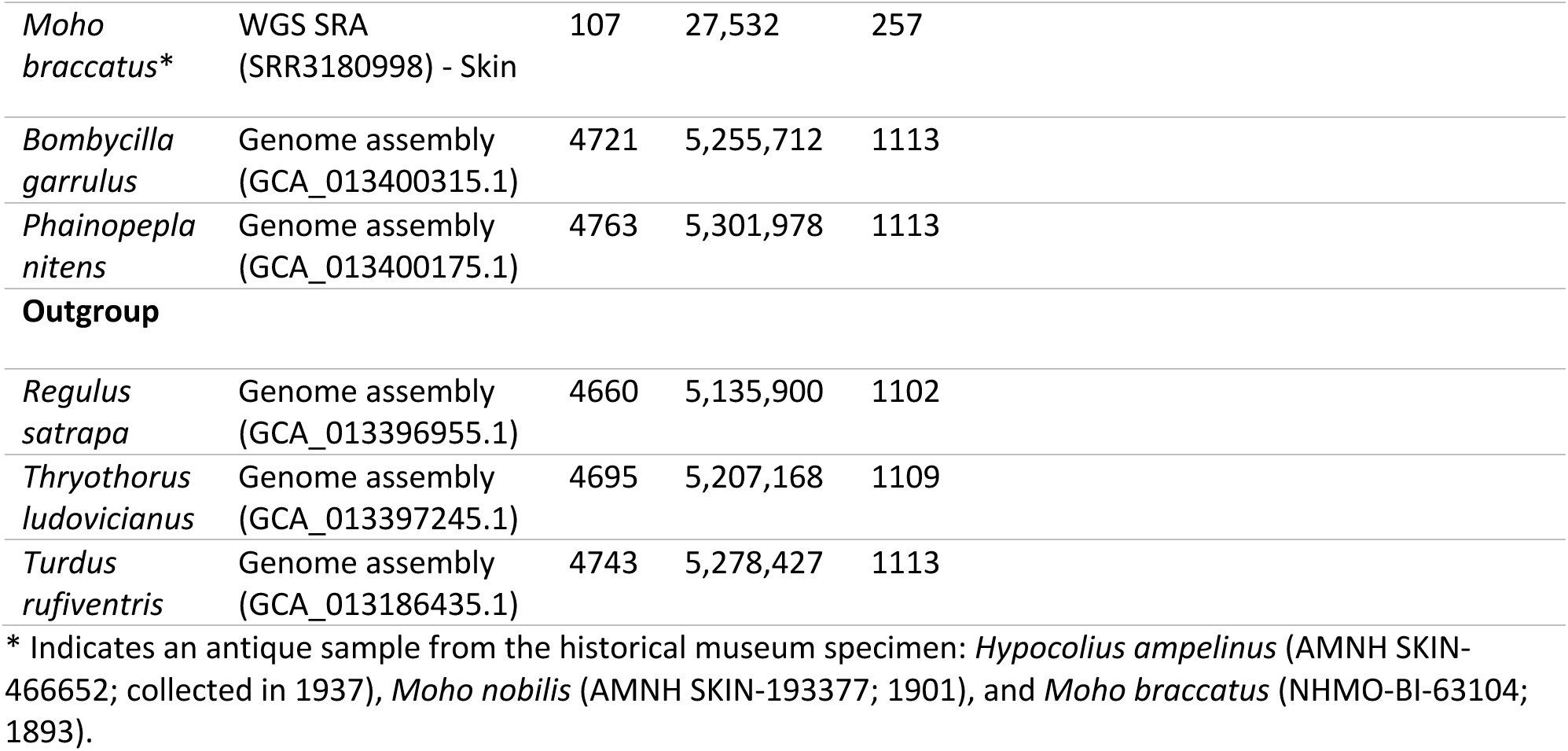
Summary statistics of UCEs extracted for each sample. For the five samples with target capture raw reads, we show statistics for both the SPAdes assembly and the Trinity assembly, as well as the number of overlapping UCEs between the two assemblies.

| Species | Data source | SPAdes assembly |  |  | Trinity assembly |  |  | Overlapping loci (before/post edge trim in PHYLUCE) |
| --- | --- | --- | --- | --- | --- | --- | --- | --- |
|  |  | Loci | Total sites | Avg. sites per locus | Loci | Total sites | Avg. sites per locus |  |
| *Dulus dominicus* | Target capture SRA (SRR2968721) | 4583 | 3,192,750 | 697 | 4513 | 3,060,487 | 678 | 4506/4505 |
| *Hylocitrea bonensis* | Target capture SRA (SRR8236567) | 4812 | 4,390,976 | 913 | 4753 | 4,216,483 | 887 | 4751/4750 |
| *Ptiliogonys caudatus* | Target capture SRA (SRR8236661) | 4827 | 4,536,869 | 940 | 4604 | 4,205,668 | 913 | 4590/4585 |
| *Hypocolius ampelinus* * | Target capture SRA (SRR8236594) - Skin | 3826 | 1,218,545 | 318 | 3075 | 885,338 | 288 | 2752/2748 |
| *Moho nobilis* * | Target capture SRA (SRR8236679) - Skin | 4307 | 1,448,938 | 336 | 3786 | 1,221,241 | 323 | 3552/3552 |
| **UCEs from Genome or WGS reads** |  |  |  |  |  |  |  |  |
|  |  | Loci | Total sites | Avg. sites per locus |  |  |  |  |

|  |  |  |  |  |
| --- | --- | --- | --- | --- |
| *Moho braccatus* * | WGS SRA (SRR3180998) - Skin | 107 | 27,532 | 257 |
| *Bombycilla garrulus* | Genome assembly (GCA_013400315.1) | 4721 | 5,255,712 | 1113 |
| *Phainopepla nitens* | Genome assembly (GCA_013400175.1) | 4763 | 5,301,978 | 1113 |
| **Outgroup** |  |  |  |  |
| *Regulus satrapa* | Genome assembly (GCA_013396955.1) | 4660 | 5,135,900 | 1102 |
| *Thryothorus ludovicianus* | Genome assembly (GCA_013397245.1) | 4695 | 5,207,168 | 1109 |
| *Turdus rufiventris* | Genome assembly (GCA_013186435.1) | 4743 | 5,278,427 | 1113 |

\* Indicates an antique sample from the historical museum specimen: *Hypocolius ampelinus* (AMNH SKIN-466652; collected in 1937), *Moho nobilis* (AMNH SKIN-193377; 1901), and *Moho braccatus* (NHMO-BI-63104; 1893).

**Table 2.** UCE datasets used in this study. All alignments underwent edge trimming in PHYLUCE unless otherwise stated.

| Number | Data set | UCE data source | Taxa | Loci | Total sites | Parsimony informative sites |
| --- | --- | --- | --- | --- | --- | --- |
| 1 | Oliveros alignment | Original alignments of Bombycilloidea and outgroup from a large supermatrix in Oliveros et al., de-gapped. | 9 | 4060 | 2,464,328 | 30,137 |
| 2 | Braun alignment | Original alignment of Bombycilloidea and outgroup from a large supermatrix in Braun et al., de-gapped. | 9 | 4307 | 2,041,232 | 20,499 |
| 3 | Oliveros alignment plus two taxa | Oliveros alignment, plus UCEs from *Phainopepla nitens* (genome) and *Moho braccatus* (WGS reads), re-aligned with the --keeplength option. | 11 | 4060 | 2,464,328 | 37,606 |
| 4 | Re-aligned Oliveros UCEs | Re-aligned raw UCEs of Bombycilloidea and outgroups from Oliveros et al. (2019) | 9 | 4820 | 2,882,679 | 41,055 |
| Re-assembled UCE data from target enrichment raw reads plus UCEs from genomes |  |  |  |  |  |  |
| 5 | SPAdes assemblies before edge trimming | All five SPAdes assemblies from UCE target enrichment reads and five genome assemblies. Without edge trimming. | 10 | 4886 | 5,741,820 | 159,462 |
| 6 | Trinity assemblies before edge trimming | All five Trinity assemblies from UCE target enrichment reads and five genome assemblies. Without edge trimming. | 10 | 4900 | 5,744,825 | 148,865 |
| 7 | SPAdes assemblies | All five SPAdes assemblies from UCE target enrichment reads and five genome assemblies. | 10 | 4886 | 4,512,375 | 104,896 |
| 8 | SPAdes assemblies + *Moho braccatus* | All above plus *Moho braccatus* | 11 | 4886 | 4,506,388 | 104,716 |
| 9 | Trinity assemblies | All five Trinity assemblies from UCE target enrichment reads and five genome assemblies. | 10 | 4900 | 4,573,118 | 104,334 |
| 10 | Mixed Trinity and SPAdes assemblies | Three Trinity assemblies for fresh tissue samples, two SPAdes assemblies for antique samples and genome assemblies | 10 | 4884 | 4,434,619 | 99,750 |
| 11 | Assembly as taxon | All five SPAdes assemblies, five Trinity assemblies, five genome assemblies, plus *Moho braccatus* | 16 | 4920 | 3,813,216 | 167,215 |
| 12 | Consensus | Consensus sequences between SPAdes and Trinity assemblies for five species, five genome assemblies, plus *Moho braccatus* | 11 | 4879 | 4,612,265 | 105,029 |

### Adding taxa to Oliveros alignment

It has been suggested that adding in taxa can improve estimates of phylogeny by “breaking up” long branches (Bergsten, 2005; Wiens and Tiu, 2012). Therefore, we added more Bombycilloidea species for which data were available to the Oliveros alignment (we chose this dataset because it was not internally trimmed). We downloaded the genome assemblies for *Phainopepla nitens* (Ptiliogonatidae) and *Bombycilla garrulus* (Bombycillidae), which were collected as part of Feng et al. (2020), from NCBI GenBank and used PHYLUCE (Faircloth, 2016) to extract UCE loci based on the 5K UCE probe set (available from https://github.com/faircloth-lab/uce-probe-sets; Sun et al., 2014). We extracted 500 bp upstream and downstream of the UCE core for each locus.

We also downloaded the WGS reads for the extinct *Moho braccatus* from NCBI SRA and used Trimmomatic v.0.39 (Bolger et al., 2014) to trim any remaining adapters. We used Bowtie2 v.2.2.5 (Langmead and Salzberg, 2012) and samtools v.1.12 (Danecek et al., 2021) to map the raw reads of *Moho braccatus* to a reference file that contained the UCEs extracted from the *Bombycilla garrulus* genome assembly. We then used SPAdes v.3.15.3 (Prjibelski et al., 2020) to assemble the mapped reads of *Moho braccatus*, with the careful flag to perform mismatch corrections, the single-cell mode (Nurk et al., 2013) to accommodate the uneven coverage of reads which is typical of target enrichment sequencing data and a coverage cutoff at five. Same as described above, we then used the standard PHYLUCE pipeline to extract UCEs from the SPAdes assembly of *Moho braccatus*.

We used MAFFT v.7.520 (Katoh and Standley, 2013) to align UCEs of *Phainopepla nitens* and *Moho braccatus* to the Oliveros alignment which were directly extracted from the original, large supermatrix as described above. To preserve the original structure of alignments, we used the add feature (Katoh and Frith, 2012) and --keeplength option in MAFFT. Since the UCEs of *Phainopepla nitens* were extracted from a genome, they are typically longer than the Oliveros alignment, therefore, we used the --addlong option of the add feature. For UCEs of *Moho braccatus* which were extracted from an antique sample, we added them in a second step using the --addfragments option. As described above, we estimated a phylogeny on data set 3 (Table 2) using IQ-TREE2 with the same settings.

### Re-aligning Oliveros UCEs

The Oliveros alignment was subsetted from a large supermatrix with over 200 taxa spanning Passeriformes, so it could include alignment errors that reflect the inclusion of distantly related taxa in the alignment; these errors may be especially problematic for taxa with poor data recovery (e.g., taxa represented by antique sample). To address this, we re-aligned the raw UCE assemblies for the six Bombycilloidea species and three outgroup species from Oliveros et al. (2019) using MAFFT with the default settings implemented in PHYLUCE. Similarly, we estimated a phylogeny using the re-aligned Oliveros UCEs (Table 2; data set 4) in IQ-TREE2.

### Re-assembling target enrichment UCE data

We also built datasets from available genome assemblies and new assemblies from target enrichment raw reads. We prioritized use of published genome assemblies over UCE target enrichment reads (Table 1). We also identified three outgroup species for Bombycilloidea that had genome assemblies available to represent Muscicapoidea, Regulidae and Certhioidea respectively.

For UCE target enrichment reads (five Bombycilloidea species), we used Trimmomatic v.0.39 (Bolger et al., 2014) to trim adapters and prinseq v.0.20.4 (Schmieder and Edwards, 2011) to filter duplicate reads. Since Oliveros et al. (2019) used Trinity to assemble fresh tissue samples and SPAdes to assemble antique samples from museum specimens, we assembled reads for all five species using both SPAdes v.3.15.3 (Prjibelski et al., 2020) and Trinity v.2.15.1 (Hölzer and Marz, 2019) to see how different assemblers may have different effects on resulting sequences. As described above, we then used PHYLUCE to extract UCEs from the SPAdes, Trinity and genome assemblies. We then performed two types of alignments: 1) using MAFFT directly without edge trimming; and 2) using MAFFT implemented in PHYLUCE where edge trimming is conducted as a default setting.

We estimated phylogenies using IQ-TREE2 on six datasets (Table 2; datasets 5-10): SPAdes and genome assemblies before and post edge trimming; SPAdes and genome assemblies with *Moho braccatus*; Trinity and genome assemblies before and post edge trimming; three Trinity assemblies for fresh tissue samples, two SPAdes assemblies for antique samples (*Hypocolius ampelinus* and *Moho nobilis*) and genome assemblies.

### Conflicting sites between SPAdes and Trinity assemblies

We evaluated base conflicts in the UCEs extracted from SPAdes and Trinity assemblies. For each of the five Bombycilloidea species, we first identified overlapping UCE loci recovered by both SPAdes and Trinity assemblies and performed pairwise alignments in MAFFT. We used snp-sites v.2.0.3 (Page et al., 2016) to generate a vcf file for each alignment and output the location of variable sites (i.e., conflicts). We counted the number of conflicting sites and tallied the type of bases in each assembly. The relative location of each conflicting site was calculated based on its position in the alignment (position divided by alignment length, where small and large numbers indicate locations near the edges and numbers close to 0.5 indicate more central location in the alignment). Since PHYLUCE performed a default edge-trimming on all alignments, we repeated this process both before and post trimming sequences.

To assess whether conflicting sites between SPAdes and Trinity assemblies would affect tree topology or branch length estimates, we conducted additional analyses in IQ-TREE2 using two datasets. First, we treated each SPAdes or Trinity assembly as a separate taxon, creating a dataset that included five SPAdes assemblies, five Trinity assemblies, five genome assemblies and *Moho braccatus* (Table 2; data set 11).

Second, we generated strict consensus sequences for the overlapping UCE loci that were recovered by both SPAdes and Trinity assemblies, using a custom Perl script; for any site where the two assemblies disagreed, we treated that site as ambiguous (“-”; equivalent to “N” or “?” in tree inference using IQ-TREE). This dataset included consensus sequences from five taxa, five genome assemblies and *Moho braccatus* (Table 2; data set 12).

### Multispecies coalescent analysis

We also estimated Bombycilloidea phylogeny by using a multispecies coalescent method, ASTRAL, to analyze the consensus UCE dataset (Table 2; data set 12). We used this dataset since it included the strict consensus sequences between SPAdes and Trinity assemblies for five species, which can minimize sequence errors for gene tree estimation. We estimated a gene tree for each locus alignment using IQ-TREE2 and combined the gene trees to estimate a species tree using weighted ASTRAL Hybrid (Zhang and Mirarab, 2022).

### Combining UCEs with legacy data

To obtain a comprehensive taxon sampling for Bombycilloidea, we combined the consensus UCE dataset with legacy data, which we define as mitochondrial sequences and nuclear gene regions used as phylogenetic markers in the pre-phylogenomic era. The legacy data available from Bombycilloidea include several mitochondrial regions, two nuclear coding regions (RAG1 and RAG2), and two nuclear introns (FIB5 and FIB7). We downloaded published mitogenomes (when they were unavailable, we downloaded individual mitochondrial regions) and nuclear loci from NCBI GenBank (accessions in Suppl. Table S1). We supplemented the sampling by extracting 13 protein-coding mitochondrial genes and two ribosomal regions from target enrichment raw reads using MitoFinder (Allio et al. 2020). To further enrich the data matrix, we extracted the four nuclear sequences available from *Bombycilla cedrorum* from the five genome assemblies (Table 1) as described by Reddy et al. (2017), using the pipeline available from https://github.com/aakanksha12/Extract_seq. We used IQ-TREE2 to estimate a phylogeny using the combined dataset (UCEs data set 12 + legacy data) as well as a phylogeny using only the legacy data.

## Results

### Phylogenetics based on different UCE data sets

In total, we estimated concatenated trees from 12 UCE datasets (Table 2). These trees exhibited two topologies which differed in the relationships among Mohoidae, Hypocoliidae and Hylocitreidae (Fig. 1). When using the original alignments extracted from the large supermatrix of Oliveros et al. (2019) and Braun et al. (2024), the trees supported a sister relationship between *Moho* and *Hypocolius*. After adding *Phainopepla nitens* and *Moho braccatus* to the Oliveros alignment and re-aligning while preserving the original alignment structure, the tree still supported that *Moho* and *Hypocolius* together were sister to *Hylocitrea*. Two datasets based on our re-assemblies of the target capture data without edging trimming also supported placing *Moho* sister to *Hypocolius*. In all of these above trees, the branch for *Hypocolius ampelinus* (based on an antique sample) was much longer than the other taxa and the other antique sample, *Moho nobilis* was also somewhat longer than the remaining taxa (Fig. 1).

However, trees estimated using the re-aligned Oliveros UCEs and datasets based on our re-assemblies with edge trimming, all supported a sister relationship between Hypocoliidae and Hylocitreidae (Fig. 1). Additionally, none of the taxa exhibited extremely long terminal branches in the trees that support a Hypocoliidae + Hylocitreidae clade.

### SPAdes vs. Trinity assemblies

When we used both assemblers to assemble the target enrichment raw reads for five species, SPAdes in general yielded more data than Trinity (both as the number of loci and the number of sites), especially for the two antique samples from museum specimens, *Moho nobilis* and *Hypocolius ampelinus* (Table 1). If the two assemblers produced similar sequences, treating each SPAdes and Trinity assembly as a separate taxon in analyses should result in very short terminal branches (little or no difference between sequences). This was true for taxa with high quality tissues but it was not true for the low-quality samples (Fig. 2a). Although the SPAdes and Trinity assemblies derived from the same taxon were consistently grouped together as “sister taxa,” the terminal branches were longer for the tips based on antique samples than for those based on fresh tissues. The tree based on this analysis supported a sister relationship between *Hypocolius* and *Hylocitrea* (Fig. 2a); i.e., it had the same topology as the trees based on methods that yielded relatively homogeneous branch lengths in Figure 1. The phylogeny built with consensus sequences between the SPAdes and Trinity assemblies exhibited the same topology and it had 100% support for placing *Hypocolius* sister to *Hylocitrea* (Fig. 2b).

**Fig. 2.**
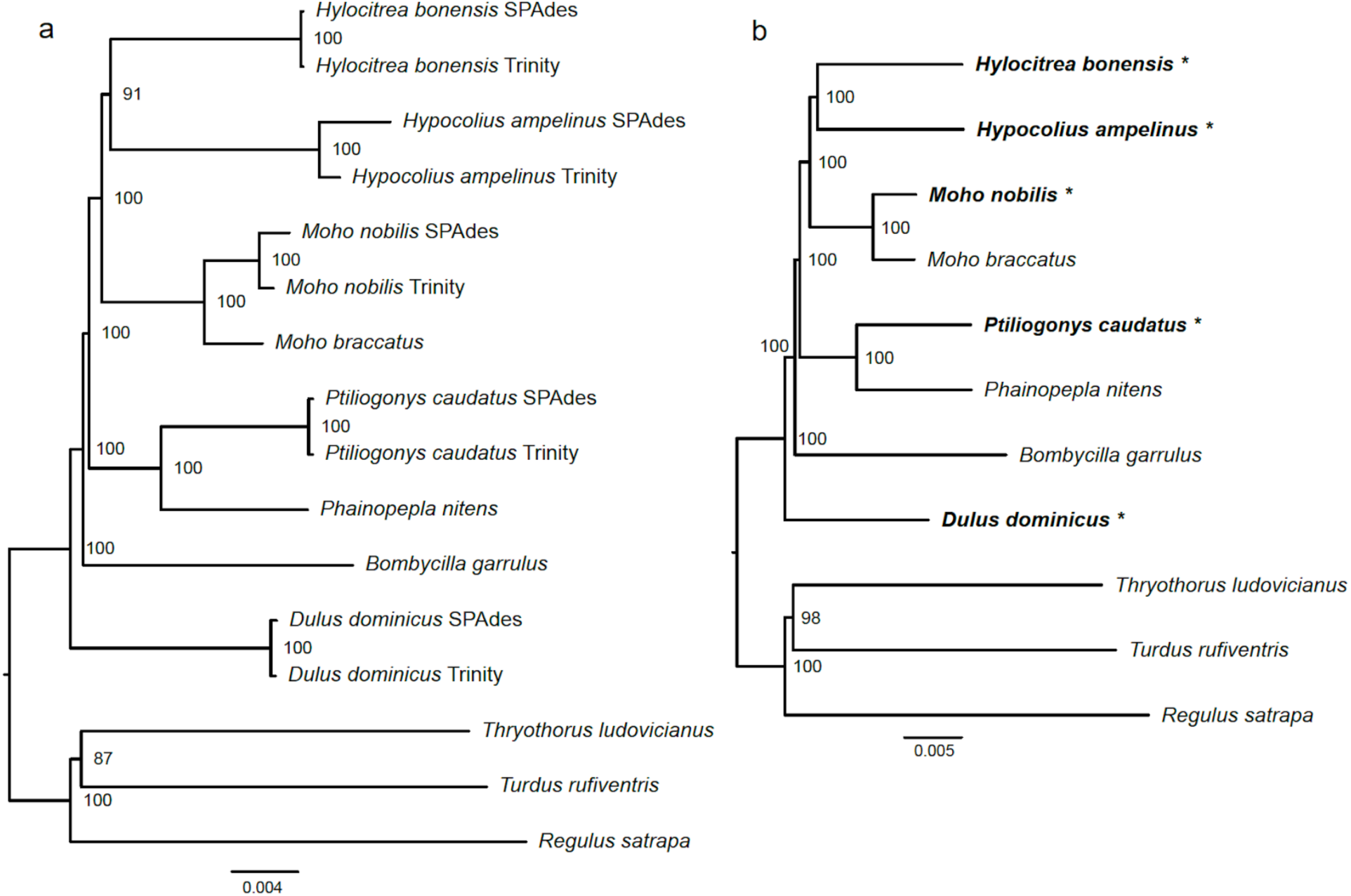
a. The maximum likelihood tree using five SPAdes assemblies, five Trinity assemblies, five genome assemblies and WGS reads of *Moho braccatus*, with each assembly treated as a taxon. b. The maximum likelihood tree estimated using the strict consensus sequences between the SPAdes and Trinity assemblies for five samples (in bold, labeled with *), five genome assemblies and WGS reads of *Moho braccatus*.

Sites that conflicted between the SPAdes and Trinity assemblies were more common at the ends of the UCE sequences (Fig. 3). For the fresh tissues, this pattern disappeared after edge trimming, but it persisted (but to a lesser degree) for the antique samples. Relative to fresh tissues, the two antique samples also had more conflicting sites in total, with proportionately more of these as sites resolved As and Ts in both assemblies (Suppl. Fig. S1). It is possible that some observed conflicts reflect heterozygous sites present in the source DNA that were resolved differently by the two assemblers, the much the larger proportion of conflicting sites in assemblies based on antique samples is more consistent with the hypothesis that most conflicts are artifactual. Transition differences (C – T and A – G) dominated the conflicts we observed for all five samples, although the proportion of transition-type conflicts was below 50% for the two antique samples, reflecting the large number of A – T conflicts.

**Fig. 3.**
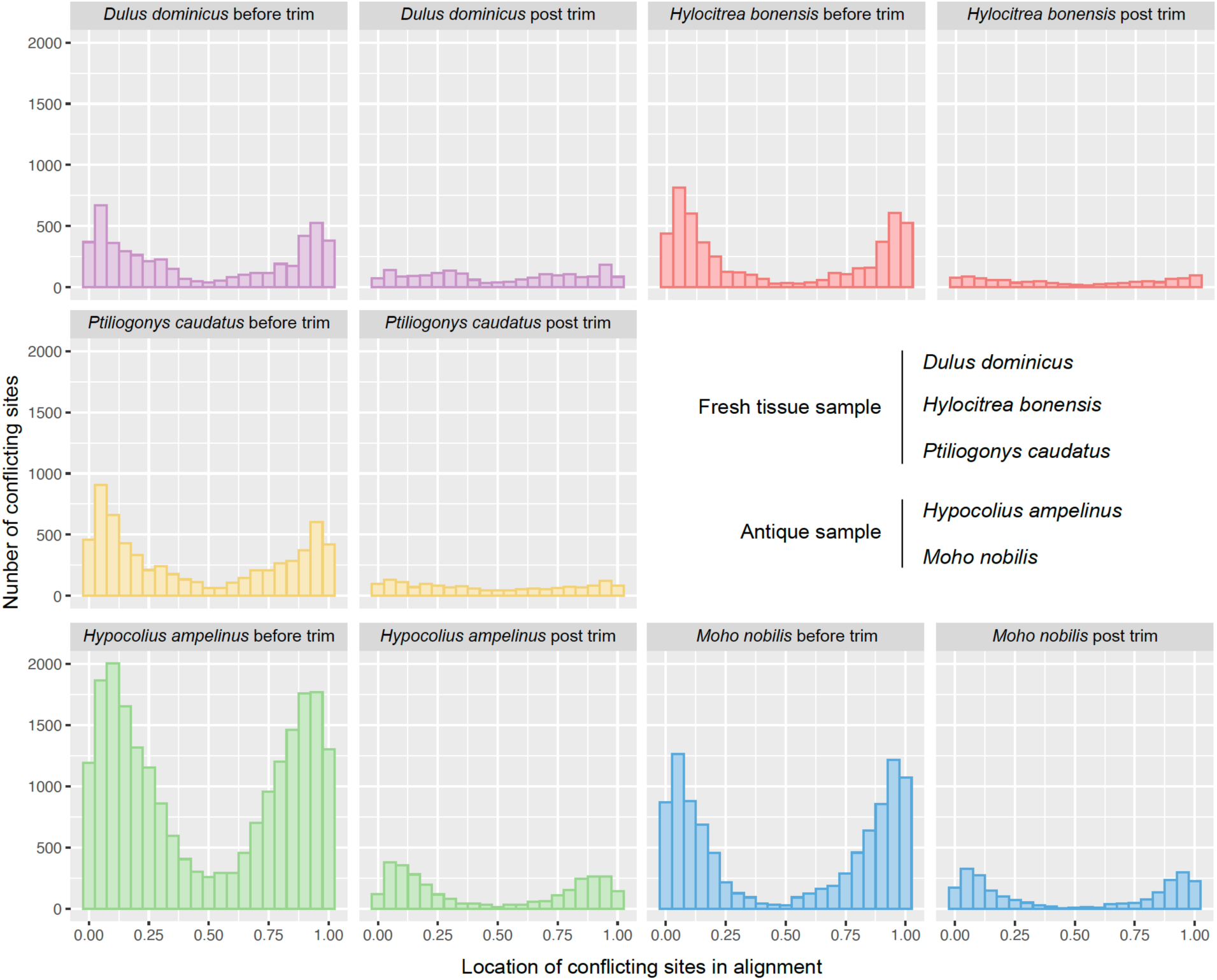
Number of conflicting sites between SPAdes and Trinity assemblies and their relative location in the alignment. From zero to one, small and large numbers indicate locations near the edges and numbers close to 0.5 indicate more central location in the alignment. For each species, we looked at alignments before and post edge-trimming.

However, we did not observe patterns that are consistent with elevated C-to-T deamination typical of damaged ancient DNA for the two antique samples.

### Multispecies coalescent method

The ASTRAL tree based on the consensus UCE dataset placed *Hypocolius* and two *Moho* species, all antique samples, at the base of Bombycilloidea as the successive sisters of the remaining members of the superfamily (Suppl. Fig. S2). The ASTRAL tree also placed *Hylocitrea* sister to *Bombycilla* with a short internal branch, conflicting with our phylogenomic trees (Figs. 1 and 2). Braun et al. (2024) explored multiple different multispecies coalescent methods and observed varying placements of the antique samples (including some outside of Bombycilloidea), but all multispecies coalescent analyses of taxa with high-quality sequences recovered the topology our concatenated analyses recover for those taxa and conflict with the topology of those taxa in our ASTRAL tree. Our ASTRAL results are consistent with both simulation and empirical studies that have suggested that gene tree summary methods tend to yield biased results for low quality samples with large amounts of missing data (e.g., Hosner et al., 2016; Xi et al., 2016; Zhao et al., 2025a). Therefore, our discussion below focused on results from the concatenation analyses.

### Comprehensive phylogeny of Bombycilloidea

Combining the consensus UCE dataset with legacy data allowed us to produce a phylogeny that included all currently recognized species within Bombycilloidea (Fig. 4). This tree shows congruent backbone topology to the trees built with re-aligned Oliveros UCEs, all datasets based on our re-assemblies with edging trimming (Fig. 1), and the consensus UCE dataset (Fig. 2). For Bombycillidae, Ptiliogonatidae, and Mohoidae, all taxa were placed within their corresponding family with strong support. The internal nodes for the clade comprising *Moho nobilis*, *M. apicalis*, and *M. bishopi* received poor bootstrap support, likely due to large amounts of missing data for the latter two taxa (which were only sampled for four and five legacy loci, respectively; Suppl. Table S1). Consistent with this hypothesis, the *Moho nobilis* + *M. apicalis* + *M. bishopi* clade had the same topology but was strongly supported when the legacy data were analyzed without UCEs (Suppl. Fig. S3; however, excluding the UCEs resulted in a different topology at the base of Bombycilloidea, with no support in that part of the tree). Regardless, the node uniting this clade and *Chaetoptila angustipluma* was strongly supported, rendering *Moho* paraphyletic.

**Fig. 4.**
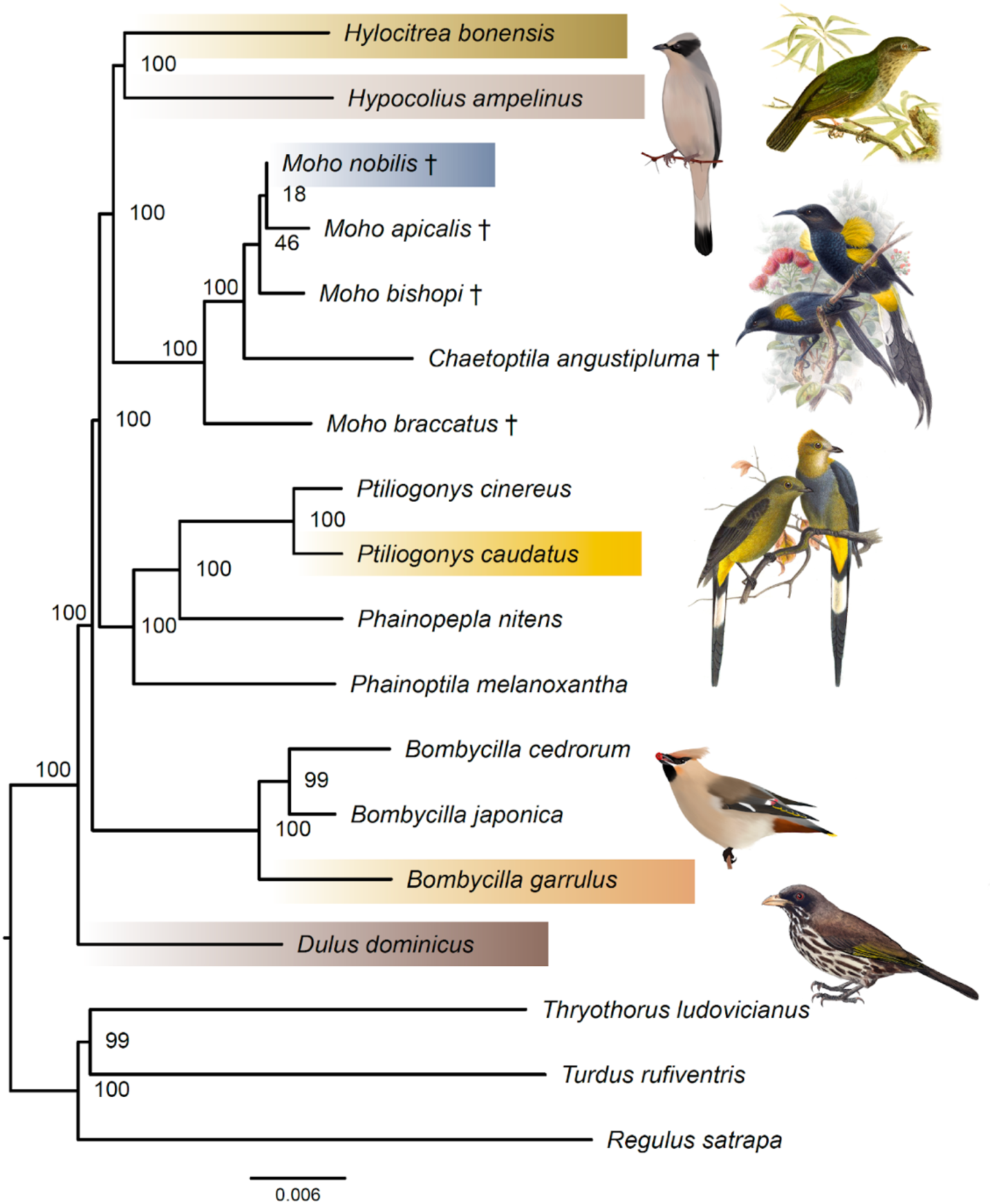
The maximum likelihood tree estimated using the consensus UCE dataset, 15 mitochondrial regions, two nuclear genes and two nuclear introns. Illustrations for *Moho nobilis* (by John Gerrard Keulemans 1893), *Ptiliogonys caudatus* (by Joseph Smit 1869) and *Hylocitrea bonensis* (from Adolf Bernhard Meyer and Lionel William Wiglesworth 1898, *The birds of Celebes and the neighbouring islands*; artist unclear) were adapted from images in Wikimedia Commons that are in the Public Domain. Illustrations of *Bombycilla garrulus* and *Hypocolius ampelinus* were drawn by M.Z. and *Dulus dominicus* was drawn by Rick Stanley.

## Discussion

Our results showed that the long branches of *Hypocolius ampelinus* and *Moho nobilis* and their sister relationship in previous work (Braun et al., 2024; Oliveros et al., 2019) were likely caused by artifactual factors such as alignment errors and missing data that reflect the quality of the source DNA. Simply adding in more taxa, e.g., *Moho braccatus*, did not break up the attraction of the two long branches. However, when the UCEs were re-aligned using a smaller, more focused taxon set, the branches were shortened and the topology shifted, resulting in a tree with a *Hypocolius* + *Hylocitrea* clade sister to *Moho*. This topology was also supported by our re-assembled datasets that were edge-trimmed and contained more loci and sites. Choice of assemblers had a large impact on the sequences recovered, especially towards the two ends of the UCEs for the antique samples. However, for our focal group, the choice of assemblers and mixed use of assemblies did not explicitly contribute to the topological conflicts observed.

### Systematics of Bombycilloidea

Bombycilloidea is part of the Muscicapida radiation which also includes Muscicapoidea, Regulidae and Certhioidea (Oliveros et al., 2019). However, compared to the latter three clades, Bombycilloidea comprises an extremely small number of species. While previous studies have reported varying relationships among the three outgroup lineages (cf. Braun et al., 2024; Oliveros et al., 2019; Stiller et al., 2024; Zhao et al., 2025b), our trees have an outgroup topology consistent with that of Stiller et al. (2024)—which utilized whole genome data—and support Regulidae as the sister group to the clade comprising Muscicapoidea and Certhioidea.

Within Bombycilloidea, our results agree with many previous studies that Dulidae is the first lineage to diverge in the superfamily (e.g., Braun et al., 2024; Burleigh et al., 2015; Oliveros et al., 2019; Spellman et al., 2008; Zhao et al., 2025b), although the exact placements of Bombycillidae and Ptiliogonatidae, as well as our focal clade (Mohoidae, Hypocoliidae and Hylocitreidae), have been more unstable. Our trees align with other genomic-scale studies (e.g., Braun et al., 2024; Oliveros et al., 2019; Zhao et al., 2025b) showing that Bombycillidae and

Ptiliogonatidae diverged successively after Dulidae, and that the clade comprising Mohoidae, Hypocoliidae, and Hylocitreidae is sister to Ptiliogonatidae. For our focal clade, our results support a sister relationship between Hypocoliidae and Hylocitreidae and together they are sister to the extinct Mohoidae.

Bombycilloidea is a species-poor but geographically widespread clade. Our phylogeny suggests an early, relatively rapid, speciation into the different families. Oliveros et al. (2019) estimated a crown age at ca. 20 million years ago (Ma) for Bombycilloidea, and the six families diverged between ca. 17–20 Ma. However, since then there has been little net diversification within any of the families. Many of the families have restricted distributions (e.g., Dulidae is endemic to the island of Hispaniola, Mohoidae were endemic to the Hawaiian Islands, and Hylocitreidae is endemic to Sulawesi), though others are much more widespread (e.g., Bombycillidae includes three waxwing species, one found throughout North America, one found in Asia, and one with a circumpolar distribution). This puzzling biogeographic pattern probably suggests a once widespread or highly vagile common ancestor (like the waxwings) that became established on various islands across the globe during the Miocene. However, it is also possible that there were pervasive extinctions in the superfamily that have obscured the biogeographic history of the group, leading to the confusing distribution that we observe today.

The common ancestor of Bombycilloidea was likely to be frugivorous given the diets of the extant species. Although the Mohoidae were primarily nectivores (leading to their initial association with the Hawaiian honeycreepers, Meliphagidae; (Lovette, 2008), they also consumed fruit (Billerman, 2020), consistent with the primarily frugivorous diets of other Bombycilloidea. Consistent with a diet primarily composed of foods that are patchily distributed in space and time (e.g., fruiting trees), and thus not readily defensible, many of these species do not defend traditional territories, with *Dulus dominicus* being highly colonial (Winkler et al., 2020a), and others, such as *Bombycilla* spp. (Winkler et al., 2020b), *Ptiliogonys caudatus*, and some *Phainopepla nitens* populations (Winkler et al. 2020c). Little is known about the behavior of the Mohoidae, but flocks outside of the breeding season has been suggested for many Mohoidae species (Billerman, 2020), as well as for many other Bombycilloidea.

The phylogeny within Mohoidae is consistent with Hennig’s “progression rule” as articulated for the Hawaiian Islands by Wagner and Funk (1995), where species colonize the oldest islands first and later disperse to younger islands. *Moho braccatus* was distributed on Kauai, the oldest of the major Hawaiian Islands (Neall and Trewick 2008), whereas all other Mohoidae (including *Chaetoptila angustipluma*) were on younger islands. Although there are exceptions to the progression rule (Heads 2010), there are many phylogenies of Hawaiian taxa that provide at least partial support for the progression rule, both in birds (Lerner et al. 2011) and in other taxa (Kleinkopf et al., 2019; Knope et al., 2020; Rubinoff and Schmitz, 2010; Rundell et al., 2004). Monophyly of *Moho nobilis*, *M. apicalis*, and *M. bishopi* is also consistent with the similarities among these taxa that were observed before their extinction (Perkins 1903); indeed, these taxa are similar enough that they have sometimes been viewed as subspecies of the same species (cf. Pratt 1979). Our phylogeny also suggests that *Chaetoptila* Sclater, PL 1871 (including *C. angustipluma* and likely the *Chaetoptila* species known from fossils) should be subsumed into *Moho* Lesson, RP 1830 based on priority of the generic names.

### Effects of inaccurate alignments and alignment trimming

Numerous studies have found that inaccurate alignments can result in erroneous topologies (e.g., Ashkenazy et al., 2019; Ogden and Rosenberg, 2006; Wang et al., 2011). Our results also reflected this. We extracted both the Oliveros and Braun alignments directly from large supermatrices that contained hundreds of taxa spanning the Avian Tree of Life and preserved their original alignment structures. These large alignments naturally included many gaps due to accumulated mutations, deletions and insertions across divergent taxa. The highly conserved UCE cores can act like anchor points during alignment to ensure that homologous sequences are easily aligned (Faircloth et al., 2012). However, since DNA from antique samples are often fragmented and damaged, UCEs extracted from these samples are usually short and contain more missing data in their flanking regions. Therefore, aligning sequences from antique samples with less closely related taxa is more likely to result in spurious alignments by chance due to their short length and gappy alignments (de Filippo et al., 2018; Landan and Graur, 2009; Smith et al., 1985). It has been suggested that large sparse supermatrices are more sensitive to phylogenetic artifacts than smaller but more complete data sets (Roure et al., 2013). To examine the role of alignment, we re-aligned the Oliveros UCEs, focusing on a smaller number of more closely related taxa and conducted edge trimming. Both re-alignment and edge trimming are likely to have reduced spuriously aligned sequences. Our results suggest that long branches can, at least in part, be driven by alignment issues when aligning divergent taxa. Thus, tackling persistent phylogenetic problems, especially those that involve large-scale genomic trees and low-quality samples, may still require examination within a more focused set of species.

Additionally, the topological discrepancy could also stem from differences in information content in addition to alignment errors, since the dataset of re-aligned Oliveros UCEs had many more loci and sites than the original Oliveros alignment (Table 2). Alignment trimming is widely used to reduce alignment errors (reviewed by Steenwyk et al., 2023), though it often comes with the cost of reducing informative sites. As implemented in the default settings of PHYLUCE, the seqcap_align program is used to trim the edges of resulting alignments to remove poorly aligned or sparsely sampled regions, ensuring that the remaining alignment has sufficient coverage across taxa and is not overly divergent from the consensus (Faircloth, 2016).

Therefore, re-aligned UCEs of the more closely related taxon set went through less aggressive edge trimming compared to the Oliveros/Braun alignments—or those that preserved the original alignment structure—which were aligned across hundreds of more distantly related taxa. The Braun alignments also underwent additional internal trimming, thus further reducing the number of sites (Table 2). However, determining the optimal amount of trimming to eliminate alignment errors without compromising phylogenetic information is not straightforward (e.g., Francis and Canfield, 2020; Portik and Wiens, 2021; Tan et al., 2015; Zhao et al., 2025b). Excessive trimming can reduce phylogenetic signal which might lead to biased phylogenies (Tan et al., 2015). For UCEs, where informative sites tend to accumulate at the two ends, UCE data can be especially sensitive to edge trimming (Portik and Wiens, 2021).

Nevertheless, failure to conduct any edge trimming resulted in exceptionally long terminal branches for two antique samples in our dataset (*Moho* and *Hypocolius*), and this is likely to have led to incorrect topologies (Fig. 1; before edge trimming).

Several studies where taxa with sequences generated from antique DNA are clustered as long branches have referred to the observed phenomenon as long branch attraction (e.g., McCullough et al. 2024; Zhao et al. 2025a), but there is a fundamental distinction between long branches due to poor sequence recovery and long branches due to accelerated sequence evolution. Susko and Roger (2021) have argued that the model space is larger for trees with long branches clustered together than trees with long branches separated and suggested this issue is central to long branch attraction; these differences in the size of model space should exist whether the basis for the long branches is biological (i.e., due to accelerated sequence evolution) or technical (i.e., reflecting the consequences of missing data in antique samples).

However, the assumption that long branch attraction can be ameliorated by using more “biologically realistic” models that better reflect the processes underlying sequence evolution (e.g., Lartillot et al., 2007; Pandey and Braun, 2020; Wang et al., 2018) clearly does not apply to long branches that reflect missing data and/or sequencing and assembly errors. It may be difficult (or even impossible) to successfully model the processes that result in long branches in trees that include antique samples. Although it is premature to propose general methods for the correction of long branches in trees with antique samples, we found that improving sequence alignments combined with edge trimming had the biggest impact in this study.

### Conflicting sites are not randomly distributed

Although choice of assemblers (SPAdes vs Trinity) did not appear to affect the topology of the Bombycilloidea tree, comparisons of the assemblies produced with the different assemblers revealed conflicting sites towards the two ends of the alignments, especially before edge trimming (Fig. 3). This partly reflects the characteristics of UCEs and target capture sequencing. UCE cores are highly conserved across vertebrates, whereas the flanking regions contain more variable sites as they locate further away from the conserved cores (Faircloth et al., 2012).

Those variable sites provide the essential information for phylogenetic inference. However, since the target capture process uses bait sets to target the desired DNA fragments (i.e., the UCE cores), and allows for selective enrichment of these regions, the flanking regions will have lower read coverage. Low read coverage is expected to reduce the ability of any assembler to detect sequencing errors and extending contigs reliably (e.g., Ayling et al., 2020; Haiminen et al., 2011).

Antique samples with DNA fragmentation and damage are likely to be prone to a higher rate of sequencing errors and the flanking regions may have even more heterogeneous read coverage. Consistent with this, we found more conflicts between the two assemblies for antique samples than the fresh tissue samples, and many conflicts persisted even after edge trimming (Fig. 3). This pattern was also reflected as relatively longer terminal branches in the tree with different assemblies treated as independent taxa (Fig. 2a). When edge trimming was used to remove potential sequencing and alignment errors, our datasets produced congruent topologies in supporting a sister relationship between *Hypocolius* and *Hylocitrea*. Both Trinity (Hölzer and Marz, 2019) and the single-cell mode of SPAdes (Nurk et al., 2013) were designed to accommodate highly non-uniform read coverage such as in transcriptome and single-cell sequencing data; while Trinity utilizes a single k-mer size when building the de Bruijn graph, SPAdes uses a set of k-mer sizes to refine the assembly with multiple steps of read error correction. Though tested with a limited number of samples, we found that SPAdes in general recovered more UCE loci and sites than Trinity, especially for antique samples (Table 1). As the true sequence is unknown, generating the consensus between the SPAdes and Trinity assemblies could be an effective way to resolve these conflicts (Fig. 2b), although this came at the cost of reducing sites since we treated conflicting sites as ambiguous. Future studies can apply these methods, e.g., treating an assembly as an independent taxon, or generating consensus sequences, to assess the performance of different assemblies and their impact on estimated phylogenies.

## Conclusions

The phylogenomic era has made it possible to imagine generating an estimate of the Tree of Life that is both complete and accurate, but it will be necessary to include data from both ancient and antique samples to achieve this goal. Antique samples may be especially important, because they can provide access not just to data from extinct taxa (like Mohoidae) and but also from extant taxa where tissues are unavailable either because they are distributed in areas without recent expeditions (like Hypocoliidae) or because their conservation status precludes their collection. Antique samples are likely to be associated with higher levels of missing data and higher rates of sequencing errors. Therefore, the sequences from antique samples may yield long branches and, potentially, incorrect topologies when they are analyzed. We suggest that studies where long branches are recovered with antique DNA samples should use multiple assembles to examine error rates, examine the impact of sequence alignment, and explore sequence trimming. In this study, these approaches allowed us to generate a strongly supported backbone phylogeny for Bombycilloidea without long branches. Combining the clean phylogenomic data with legacy sequence data ultimately allowed us to generate a complete phylogeny for the superfamily.

## Data availability

The data that support the findings of this study (alignments and tree files) are openly available in figshare at http://doi.org/10.6084/m9.figshare.28255466.

## Acknowledgements

We thank Rick Stanley for drawing the palmchat illustration. The Kimball-Braun lab provided helpful comments on earlier drafts of this manuscript. This work was supported by a National Science Foundation grant (DEB 1655683) to RTK and ELB.

