## Supplementary material for "The phylogenetic position of the extinct Hawaiian honeyeaters: Overcoming the limitations of antique DNA": Suppl.

**Table S1**. GenBank accessions for legacy loci. For nuclear loci marked with “extract”, they were extracted from genome assemblies. For mitochondrial region marked with “1”, they were extracted from published mitogenomes or target enrichment raw reads using MitoFinder.

| **Species** | **RAG1** | **RAG2** | **FIB5** | **FIB7** | **mitogenome** | **ATP6** | **ATP8** | **CO1** | **CO2** | **CO3** |
|---|---|---|---|---|---|---|---|---|---|---|
| *Bombycilla cedrorum* | FJ177356 | FJ177348 | EF468341 | EF471862 | KJ909187 | 1 | 1 | 1 | 1 | 1 |
| *Bombycilla garrulus* | extract | extract | extract | extract | SRR2968728 | 1 | 1 | 1 | 1 | 1 |
| *Bombycilla japonica* | FJ177357 | FJ177349 | | | | | | KF946605 | | |
| *Chaetoptila angustipluma* | FJ378045 | | | | | | | | | |
| *Dulus dominicus* | AY319980 | FJ177350 | | | SRR2968721 | 1 | 1 | 1 | 1 | 1 |
| *Hylocitrea bonensis* | FJ177360 | FJ177355 | | | SRR8236567 | 1 | 1 | 1 | 1 | 1 |
| *Hypocolius ampelinus* | FJ177361 | | | | SRR8236594 | | | 1 | | 1 |
| *Moho apicalis* | FJ378046 | | FJ378049 | | | | | | | |
| *Moho bishopi* | FJ378042 | | FJ378047 | | | FJ383119 | | | | |
| *Moho braccatus* | | | | | NC_031348.1 | 1 | 1 | 1 | 1 | 1 |
| *Moho nobilis* | FJ378066 | | FJ378051 | FJ378053 | SRR8236679 | | 1 | 1 | | 1 |
| *Phainopepla nitens* | AY319995 | FJ177351 | extract | extract | NC_053061.1 | 1 | 1 | 1 | 1 | 1 |
| *Phainoptila melanoxantha* | AY307204 | FJ177352 | | | | AF273918 | AF273918 | JQ175765 | | |
| *Ptiliogonys caudatus* | FJ177358 | FJ177353 | | | SRR8236661 | 1 | 1 | 1 | 1 | 1 |
| *Ptiliogonys cinereus* | AY443324 | AY443215 | | | | | | | | |
| **Outgroup** | | | | | | | | | | |
| *Regulus satrapa* | AY443327 | AY443221 | extract | extract | NC_051014 | 1 | 1 | 1 | 1 | 1 |
| *Thryothorus ludovicianus* | MG495479 | MG495505 | extract | extract | NC_051032 | 1 | 1 | 1 | 1 | 1 |
| *Turdus rufiventris* | extract | extract | extract | extract | NC_028179 | 1 | 1 | 1 | 1 | 1 |

**Table S1** continue.

| **Species** | **Cytb** | **ND1** | **ND2** | **ND3** | **ND4** | **ND4L** | **ND5** | **ND6** | **rrnL** | **rrnS** |
|---|---|---|---|---|---|---|---|---|---|---|
| *Bombycilla cedrorum* | 1 | 1 | 1 | 1 | 1 | 1 | 1 | 1 | 1 | 1 |
| *Bombycilla garrulus* | 1 | 1 | 1 | 1 | 1 | 1 | 1 | 1 | 1 | 1 |
| *Bombycilla japonica* | AF285797 | | FJ177332 | | | | | | AF151411 | |
| *Chaetoptila angustipluma* | FJ383122 | | | | | | | | | FJ378061 |
| *Dulus dominicus* | 1 | 1 | 1 | 1 | 1 | 1 | 1 | 1 | 1 | 1 |
| *Hylocitrea bonensis* | 1 | 1 | 1 | 1 | 1 | 1 | 1 | 1 | 1 | 1 |
| *Hypocolius ampelinus* | 1 | 1 | | 1 | 1 | 1 | | 1 | | |
| *Moho apicalis* | FJ383123 | | | | | | | | | FJ378058 |
| *Moho bishopi* | FJ383124 | | | | | | | | | FJ378059 |
| *Moho braccatus* | 1 | 1 | 1 | 1 | 1 | 1 | 1 | 1 | 1 | 1 |
| *Moho nobilis* | | 1 | | 1 | 1 | | 1 | 1 | | 1 |
| *Phainopepla nitens* | 1 | 1 | 1 | 1 | 1 | 1 | 1 | 1 | 1 | 1 |
| *Phainoptila melanoxantha* | FJ177323 | | FJ177338 | | | | | | | |
| *Ptiliogonys caudatus* | 1 | 1 | 1 | 1 | 1 | 1 | 1 | | 1 | 1 |
| *Ptiliogonys cinereus* | AY443258 | | FJ177341 | | | | | | | |
| **Outgroup** | | | | | | | | | | |
| *Regulus satrapa* | 1 | 1 | 1 | 1 | 1 | 1 | 1 | 1 | 1 | 1 |
| *Thryothorus ludovicianus* | 1 | 1 | 1 | 1 | 1 | 1 | 1 | 1 | 1 | 1 |
| *Turdus rufiventris* | 1 | 1 | 1 | 1 | 1 | 1 | 1 | 1 | 1 | 1 |

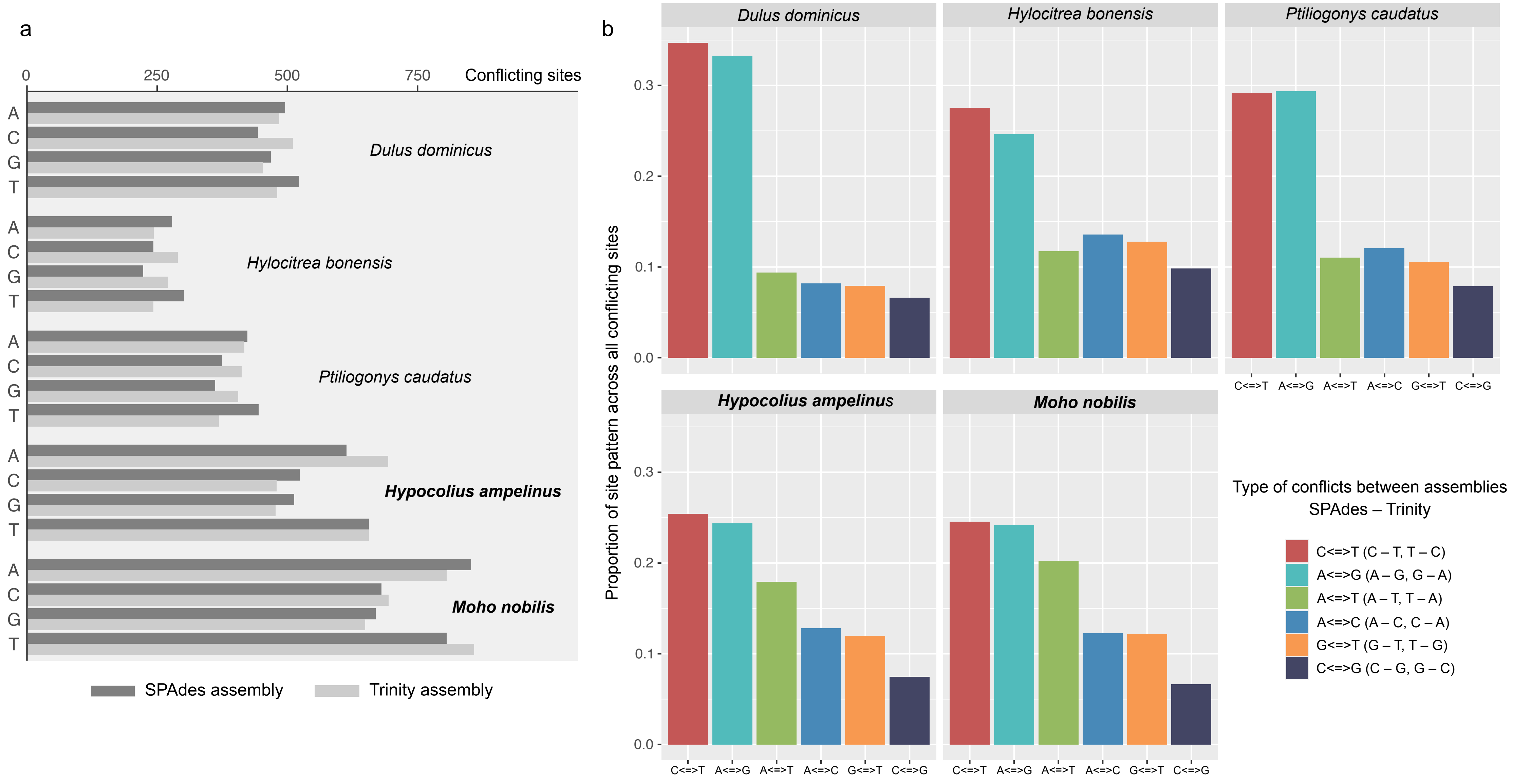


**Fig. S1**. Base composition of conflicting sites between post-trim SPAdes and Trinity assemblies. a. The number of each base summarized from all conflicting sites. b. Proportion of each site pattern across all conflicting sites between the two assemblies. For example, A <=> T indicates a site where SPAdes recovers as A while Trinity recovers as T (A – T) or SPAdes recovers as T while Trinity recovers as A (T – A).

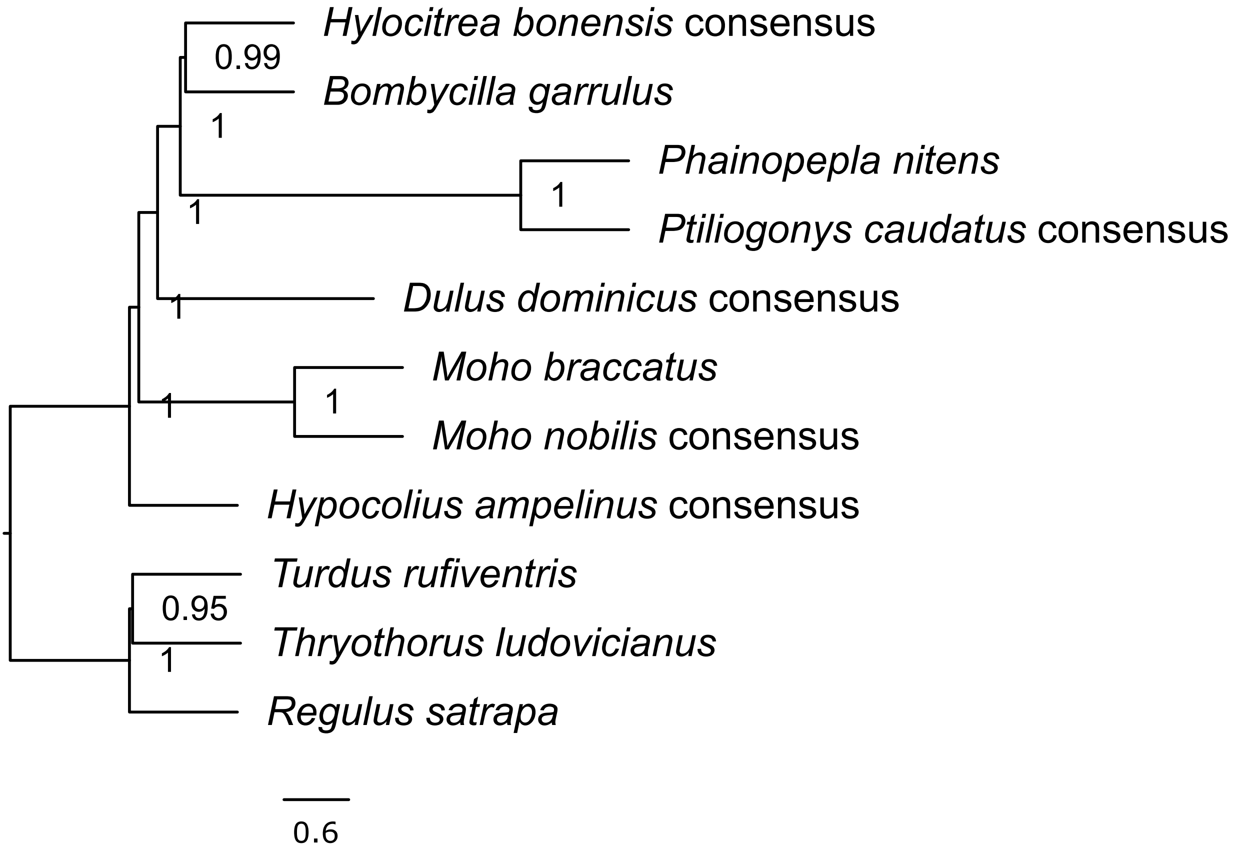


**Fig. S2**. Weighted ASTRAL built using gene trees estimated using the consensus UCE dataset. Internal branch lengths are coalesent units and terminal branch lengths are arbitrary. Support values are local posterior probabilities.

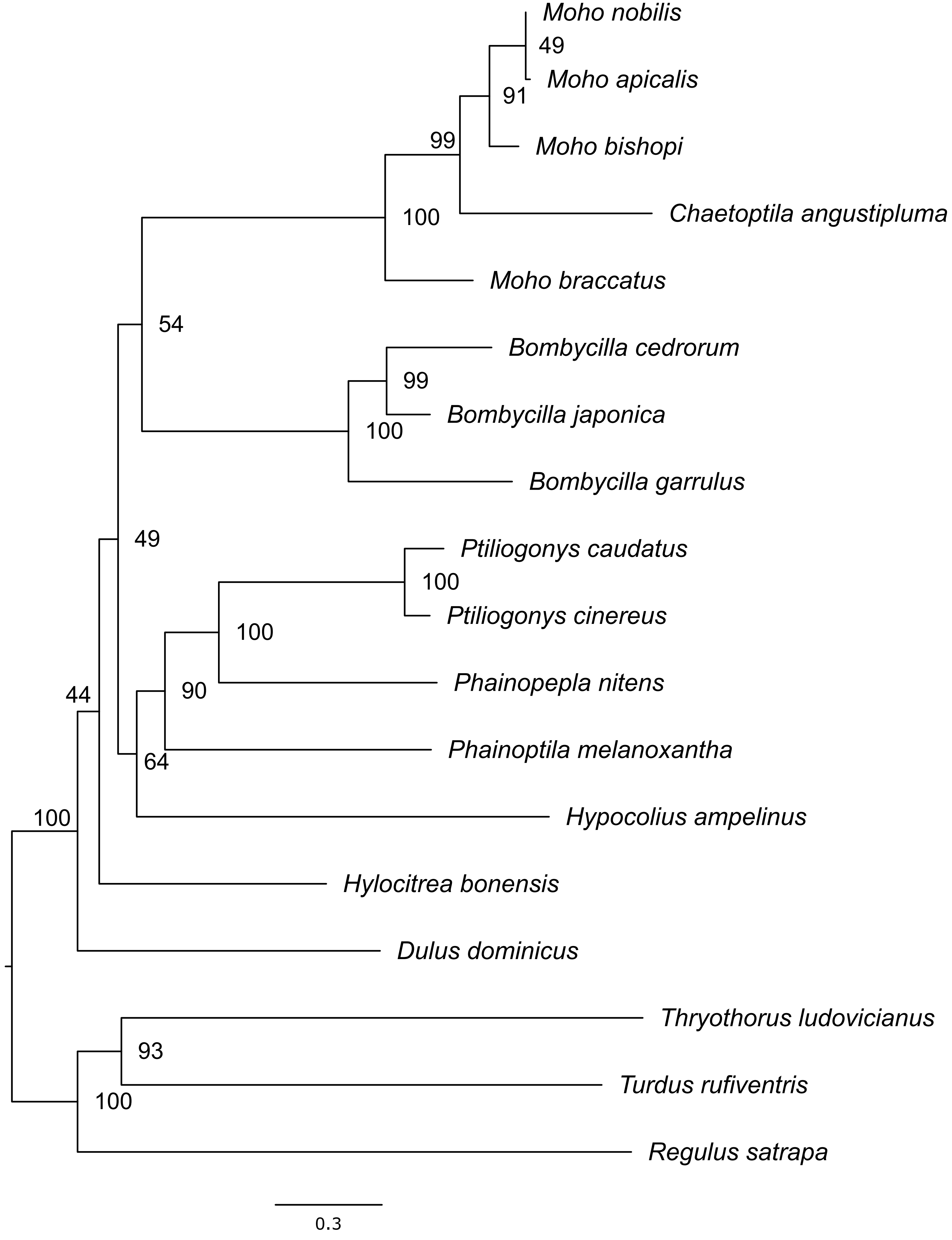


**Fig. S3**. Maximum likelihood tree estimated based on the concatenated legacy loci (mitochondrial and nuclear loci combined) using IQ-TREE2. Values at nodes are ultrafast bootstrap support values based on 1000 replicates.
